# Widespread changes in firing rate and functional connectivity across the fronto-parietal network during rule guided working memory

**DOI:** 10.1101/479410

**Authors:** RF Salazar

**Affiliations:** Department of Basic Neuroscience, University of Geneva, Switzerland.

**Keywords:** Cognitive set-switching, prefrontal cortex, posterior parietal cortex

## Abstract

Flexible rule-based behavior is an integral component of our daily lives. Rule dependent changes in the activity and functional connectivity of neurons in the frontoparietal network may underlie this flexibility. To test this idea, we simultaneously recorded neural activity from multiple areas in prefrontal and posterior parietal cortices of macaque monkeys while they performed a delayed match-to-sample task involving a non-instructed switch between location and identity matching rules. Our analysis revealed rule-dependent differences in firing rates during all phases of the task and marked task-dependent increases in spike count correlations with only weak differences between rules. These effects were widespread with a high incidence occurring within and between the dorsolateral prefrontal cortex, the lateral intraparietal area and area PG. We conclude that rule based visual working memory is associated with widespread modifications in excitability and functional connectivity across the frontoparietal network.

---

Daily life requires the continuous integration of sensory information with internally generated plans to complete a nearly constant stream of goal-oriented behaviors. Often, environmental information must be used in different ways depending on context, requiring the generation of internal guidelines, or ‘rules of the game’ (Miller, 2000). Once formed, these rules, representing stable relationships between context and behavior, can be deployed at will. Deficits in the flexible deployment of behavioral rules have been repeatedly found in individuals suffering from certain psychiatric illnesses, such as schizophrenia (Lesh et al., 2011), which makes understanding the mechanisms underlying rule guided behavior a high priority.

The neuronal signals, which guide flexible rule-based behavior, are closely related to the “top-down” signals thought to control attention (Kastner and Ungerleider, 2000; Gazzaley et al., 2007; Bressler et al., 2008), and are expressed as task-related changes in activity recorded in widespread regions of prefrontal and posterior parietal cortices (Johnston et al., 2007; Stoet and Snyder, 2009; Goodwin et al., 2012; Crowe et al., 2013). Similarly, working memory processes are supported by distributed patterns of neuronal activity and functional interactions encompassing multiple cortical areas across the fronto-parietal network (Hebb, 1949; Pesaran et al., 2002; Siegel et al., 2009; Fell and Axmacher, 2011; Fuster and Bressler, 2012; Salazar et al., 2012; Roux and Uhlhaas, 2014; Dotson et al., 2014). However, the contributions of large-scale functional interactions to rule guided working memory processes are still largely unknown.

Here, we analyzed the neuronal activity (firing rates) and functional connectivity (trial-to-trial spike count correlations) within and between areas of the prefrontal and posterior parietal cortices from monkeys performing a rule-based, delayed match-to-sample task. By comparing these quantities between different matching rules, we addressed the following questions: 1) What is the distribution of rule-selective neurons in the fronto-parietal network? 2) How does the incidence, magnitude, task, and rule dependence of correlated firing vary across the fronto-parietal network?

## MATERIALS AND METHODS

### Behavioral Paradigm and Performance Criterion

The task consisted of an oculomotor delayed match-to-sample task involving a non-instructed rule switch (Salazar et al., 2012) (Fig. 1a). Briefly, the monkeys (*Macacca mulatta*) had to fixate a central dot while a sample object was presented at one out of three possible locations. After a random delay (800 to 1200 ms), a match stimulus, consisting of two objects (target and distractor), was presented at two of the three possible locations. Correct responses required a saccade to the object matching either the identity (identity rule) or location (location rule) of the sample object. Behavioral performance was calculated using a 100 trial sliding window with a 1 trial step (Fig. 1b,c). After 300 consecutive windows were above 80%, the rule in effect was changed without cueing the animal. Special care was taken to alternate the daily order of the starting rule to balance the sequence of the task. Several switches between rules could occur each day, but most often a single successful switch was observed. Typically, the behavioral criterion on the 1^st^ rule was reached within 300-500 trials.

**Figure 1.**
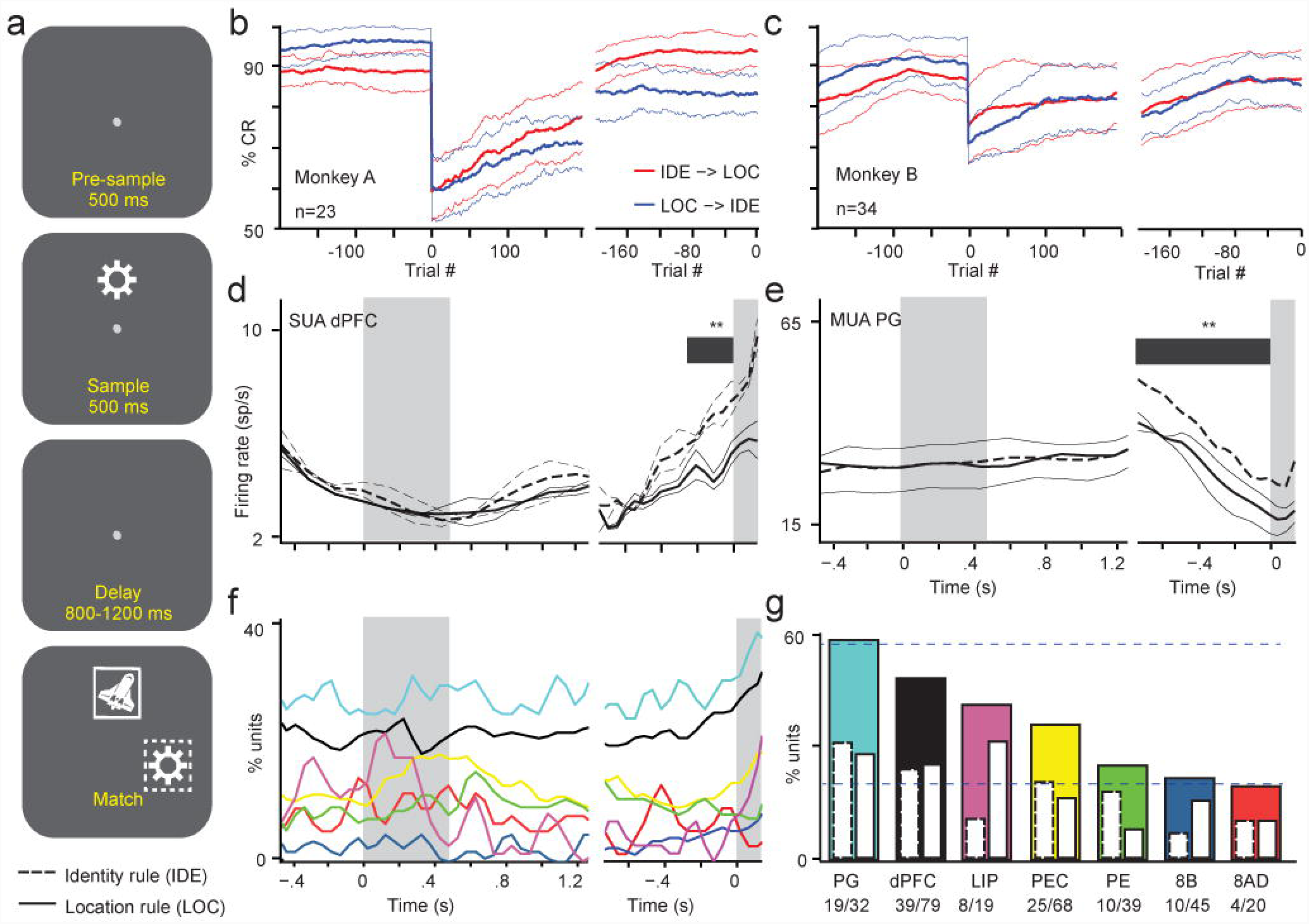
Rule specific activity in prefrontal and posterior parietal neurons. (a) Schematic drawing illustrating the behavioral task. (b, c) Behavioral performance is shown for Monkey A and Monkey B, respectively. The performance of the animals was calculated using a 100 trial window stepped by one trial until a rule switch occurred (Trial # 0). After the first switch, the next 100 trials were used to calculate the performance on trial 1 of the second rule and the same stepping procedure was applied. The subplots on the right correspond to the trials before the behavioral criterion was reached on the 2^nd^ rule (Trial #0). The performance over all sessions (mean ± std) was calculated separately for switches occurring from the identity to the location-matching rule (red: IDE -> LOC) and vice versa (blue: LOC -> IDE). (d, e) Two example peri-stimulus time histograms (PSTH) are shown for a single-unit recording in dPFC (d) and a multi-unit recording in area PG (e). The left and right portions of the plots are locked to the sample onset and match onset, respectively. Plots were smoothed using a 2 bin moving average (solid lines: location rule; dashed lines: identity rule). When multiple blocks for one rule occurred, the average PSTH across blocks (thick lines) and the PSTHs for the individual blocks (thin lines) were calculated. When only one block for a rule occurred then only a single thick line was used (e). Dark grey bars indicate significant differences between the two rules (p<0.01, Wilcoxon Rank Sum test). Shaded grey areas indicate the sample and match periods of the task. (f, g) The incidence of rule selective units is shown as a function of time (f) and cortical area (g). The same colors are used in (f) and (g) for each area. The inset bars in (g) indicate the preferred rule (solid lines: location rule; dashed lines: identity rule). The number of significant units with respect to the sample size for each area is given below the cortical area abbreviations. Horizontal dashed blue lines show the confidence intervals, 0.05% and 99.95%, for a random and uniform distribution of rule selective units.

The animal’s performance would often fluctuate throughout recording sessions, and would dip during rule switches. In order to ensure that we only analyzed data during time periods when the animal’s performance was stable, we implemented several procedures for trial selection. To identify when performance became stable after a rule switch, we removed the first 25 trials and trials with reaction times below 80 ms, and then calculated the performance using the rest of the trials. If the performance was below 75%, the first remaining trial was iteratively removed until performance was ≥ 75%. In addition, we used a sliding window of 50 trials stepped one trial at a time to create a performance curve used to identify short drops in performance. When the minimum value of this curve was below 55%, the block of 50 trials used to calculate this minimum value was excluded and the performance curve was calculated again. These steps were repeated until no value in the curve was below 55%. Drops in performance were observed only in Monkey B in 11 out of 34 sessions. In eight sessions, the removal of one block of 50 trials was sufficient to have the performance curve above 55%. In the remaining three sessions, two, three and five blocks of 50 trials were removed.

### Neurophysiological Recordings and Data Processing

We recorded broadband neuronal activity (1 Hz – 10 kHz, sampled at 30 kHz) from prefrontal and posterior parietal cortices in two monkeys (Monkey A and B). Acute recordings were performed in Monkeys A and B and semi-chronic recordings were later performed in a second round of recordings in Monkey B (for more details see Salazar et al., 2012; Dotson et al., 2015). All procedures involving the animals were performed in accordance with NIH guidelines and the Institutional Animal Care and Use Committee of Montana State University. Areal designations of the recording sites were determined either by the site of electrode penetrations combined with depth readings or by histological sectioning (for more details see Salazar et al., 2012). Spike waveforms were extracted from the highpass-filtered signal (500 Hz – 7 kHz) on each electrode using an extraction threshold of 4 (acute recordings) or 5 (semi-chronic recordings) standard deviations of the background noise, calculated from the whole recording session. Spike sorting was performed using the MClust software package (http://redishlab.neuroscience.umn.edu/MClust/MClust.html) and artifacts were removed by visual inspection. Single units were considered for further analyses only if they were isolated during the whole recording session and displayed a clear refractory period in their inter-spike interval histogram. Multiunit recordings displaying evidence of instability (large variations in amplitude over time) were excluded from the analyses.

### Firing Rate Analyses

To test for differences in neuronal firing rates between the two rules, the spike trains were pooled across stimuli and binned at 50 ms intervals (no overlap). Because the duration of the delay period varied across trials, the spike trains were aligned and referenced either to the sample onset occurring at 0 ms or to the match onset which occurred randomly between 800-1200 ms following the offset of the sample. In the sample onset alignment, the tested period for statistics lasted 1800 ms (pre-sample, sample and delay) whereas in the match onset alignment, the tested period lasted 800 ms (delay only). The mutual information (MI) between the resulting spike-counts and the rule was calculated using a MI toolbox (Peng et al., 2005) and tested for significance at each time-bin. Significance was assessed using the probability distribution of a surrogate population. This population was created by shuffling (500 iterations) the rule assignment of each trial while maintaining the same total number of trials in each rule. A unit was considered to have a different firing rate between the two rules, referred to as rule-selective unit, if at least two time bins were significant (p < 0.026 with Bonferroni correction) during the sample-aligned time series (1800 ms) or at least one time bin during the match-aligned time series (800 ms). Rule preference was simply assigned to the rule with the higher firing rate.

### Spike-Count Correlation Analyses

Spike count correlations (r_sc_) were calculated by finding the Pearson correlation coefficient between spike counts across trials. Practically, we applied the MATLAB function “corrcoef” to the numbers of spikes of two units (i and j). This function uses the covariance C and the following formula:

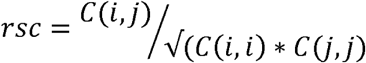

This quantity has also been referred to as the noise correlation calculated directly from spike counts (Bair et al. 2001). To control for different experimental conditions (e.g. stimulus location/identity, behavioral rule), we normalized each r_sc_ value by subtracting its mean. To test for significance and to control for the influence of firing rate on these correlations, we created a surrogate population by randomizing the trial sequence (500 iterations). Because the number of iterations limits the lowest accessible p-value and because the calculations are CPU-time consuming, we used a fitting method to estimate lower p-values. We fit a generalized extreme value distribution (GEV) to the probability density function of the surrogate distribution (Figure S1). The GEV distribution served as a look up table for p-values. In this way, we were able to estimate p-values beyond those available from the surrogate distribution alone. We chose a two-sided false discovery rate of .01 (negative and positive correlations). These criteria were applied to four 500 ms epochs for each rule. These epochs included the period prior to sample onset (pre-sample), the full duration of the sample period (sample), the early delay immediately following the offset of the sample (delay 1), and the late delay prior to match onset (delay 2). A pair of units was considered as having significant correlated firing if the correlation coefficient was significant in at least one of the four epochs on one of the rules.

### Coherence Analysis

To evaluate the correlations between pairs of units as a function of frequency, we calculated the spectral coherence between spike trains using a point-process multi-taper analysis (Mitra and Pesaran, 1999) implemented in the Chronux toolbox (Bokil et al., 2010) with three tapers and a time-bandwidth product of two. The test for significance was applied separately for each epoch of the task using a surrogate distribution of coherence values computed on the trial-shuffled spike trains (n = trial count - 1). The 99^th^ percentile of this distribution was used as a threshold, corresponding to a p-value of 0.01. Because the spectral resolution depends on the number of spikes, it was necessary to interpolate the spectra at 1 Hz, using a linear estimation to combine the results from different pairs of units.

## RESULTS

We recorded neuronal activity simultaneously from multiple prefrontal and posterior parietal areas of two monkeys (Monkeys A and B) while they performed an oculomotor, delayed match-to-sample task incorporating a non-instructed switch between location- and identity-matching rules (Fig. 1a). Correct responses required the animal to make a saccade, after a variable delay period (800 – 1200ms), to the object matching either the identity (identity rule) or location (location rule) of the sample stimulus. Importantly, the visual information was the same for both rules, thus we could assess the influence of endogenous processes on neuronal activity. Trials for each rule were presented in blocks and the matching rule was switched without cueing the animal after criterion performance was reached (>80% correct responses for 300 trials). Monkeys A and B successfully switched between the two rules in 23 and 34 daily sessions, respectively. Figure 1b and 1c show, respectively, the average behavioral performance for the two animals during a rule switch and prior to reaching stable performance on the second rule. To ensure an accurate estimate of spike-count correlations, only units (multi- or single-unit) having an average firing rate above 2 Hz during the memory delay of both matching rules were included in this analysis (Fig. S2). Also, only cortical areas with at least 15 units that passed the firing rate criteria were considered in this study. Using these criteria, a total of 302 units (277 multi-units and 25 single-units were pooled together and referred to as units; 17% of these units originated from Monkey A and 14%/69% from Monkey B during acute/chronic recordings) from three prefrontal areas (8AD, 8B and dPFC) and four posterior parietal areas (PE, PEC, LIP and PG), were used in this study (see Materials and Methods for details). During semi-chronic recordings in Monkey B, not all electrodes were moved between sessions; subsequently, 36% of the units and 18% of the pairs of units were recorded from electrodes at the same sites.

### Rule Specificity in Firing Rates

To determine the rule dependence of neuronal responses, we calculated the firing rate of units on correct trials (across all stimuli) during periods of stable behavioral performance and then tested for time-dependent differences in firing rate between the identity- and the location-matching rules. We found rule-dependent differences during the memory delay and all other periods of the task (Fig. 1d and e), including the period before the sample onset. These effects are illustrated by the incidence of significant rule-dependent differences in firing rate across the population of units in each area as a function of time (Fig. 1f). Some cortical areas displayed increased rule-dependent activity during the sample period (e.g., PEC and LIP) or prior to the match period (e.g., PG and dPFC) while other areas displayed relatively little time-dependent variations across the population (e.g., 8AD and 8B). We also found different proportions of rule-selective units across areas. Consistent with previous reports (Wallis et al., 2001; Stoet and Snyder, 2004), dPFC and areas lateral to or within the lateral bank of the intra-parietal sulcus (PG, LIP) had the highest proportion of rule selective units (Fig. 1g); notably, area PEC also displayed a relatively high proportion (37%) of rule selective units. The proportions of rule selective units differed significantly between the different cortical areas (dashed horizontal lines; p<0.01 two-sided shuffled surrogates). We identified the preferred rule for each rule-selective unit in each cortical area by selecting the rule in which the firing rate was the highest (inset bars Fig. 1g). Similar to a previous report on prefrontal cortex (White and Wise, 1999), some units preferred the identity rule while others preferred the location rule. The respective proportions of these units varied among the different cortical areas (e.g., LIP) but, across the whole population, we found no significant difference in the incidence of rule preference (p>0.42, two-sample Kolmogorov-Smirnov test).

### Functional Connectivity Analysis

While these findings demonstrate widespread rule specific activity throughout the task, how populations of neurons cooperate in large-scale circuits to implement rule-guided behavior is largely unknown. Therefore, we examined the task dependence of functional connectivity by calculating spike count correlations (r_sc_) (Cohen and Newsome, 2008) as previously described (Zohary et al., 1994; Lee et al., 1998; Constantinidis and Goldman-Rakic, 2002; Cohen and Newsome, 2008). Before determining if the patterns of functional connectivity differed between rules, we first evaluated the task-dependence of r_sc_ for the two rules separately. Using only correct trials during periods of stable behavioral performance, r_sc_ was calculated for all pairs of units during four 500 ms epochs aligned to different events of the task (pre-sample, sample, delay 1, and delay 2). To avoid any potential cross talk between neuronal signals (Ecker et al., 2010), only pairs of units recorded on different electrodes (n=1430) were considered. We refer to pairs of units with at least one epoch having a significant r_sc_ (false discovery rate < 0.01, two-sided, shuffled surrogates) during either rule as functionally connected units. This analysis revealed that 24% of the pairs of units were positively correlated (Fig. 2 in red) and 10% were negatively correlated (Fig. 2 in green), confirming previous reports that positive correlations are generally more prevalent (Zohary et al., 1994; Constantinidis and Goldman-Rakic, 2002; Cohen and Maunsell, 2009). Because the percentage of positive r_sc_ values was much higher, we focused the following analyses on them. During the sample, delay 1 and delay 2 epochs, the incidence of significant positive correlations increased, respectively, by 150%, 154% and 140% compared to the pre-sample epoch. These increases were not due to changes in firing rates because the shuffled surrogates were calculated independently for each epoch. Thus, functional connectivity is modulated by different events of the working memory task.

**Figure 2:**
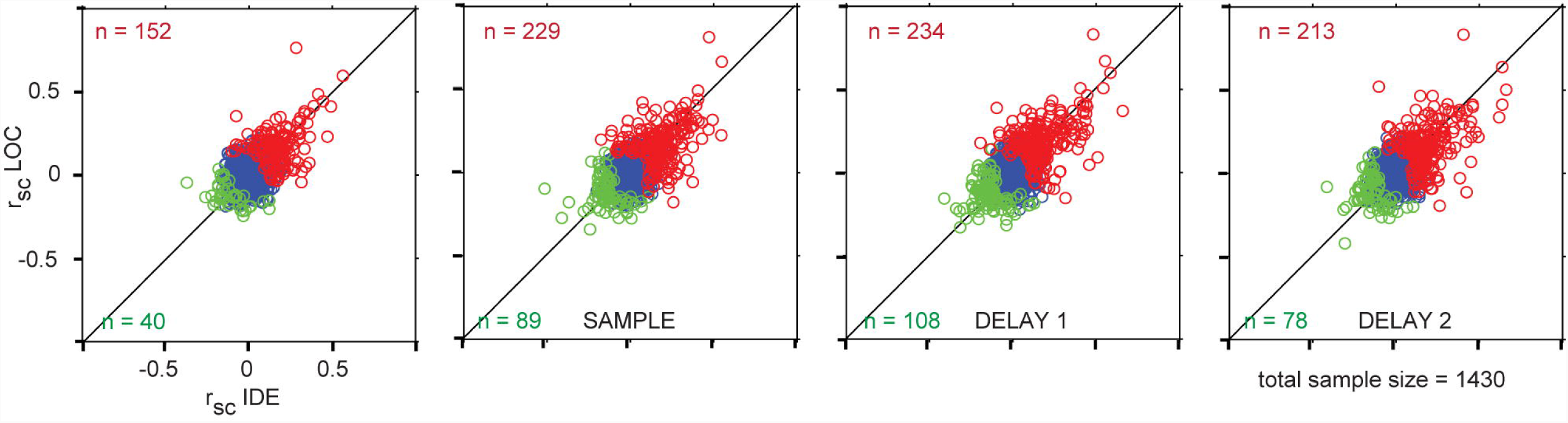
Task-dependent spike-count correlations in the fronto-parietal network. Scatter plots of the spike-count correlations (r_sc_) during the identity (x-axis) and location (y-axis) rules for all pairs of units. Each panel corresponds to a different period of the task. From left to right, presample (500 ms window prior to sample onset), sample (500 ms window prior to sample offset), delay 1 (500 ms window after sample offset), and delay 2 (500 ms window prior to match onset). The color of the circles corresponds to negative correlations (green), positive correlations (red) or non-significant correlations (blue). The numbers of pairs with significant positive or negative correlations for at least one rule are shown in red (top of plot) and green (bottom of plot), respectively.

To determine the influence of the matching rule on functional connections, we used two different approaches: one at the population level and one at the pairwise level. First, we tested whether the distribution of differences in r_sc_ between the two rules (IDE r_sc_ - LOC r_sc_) differed from zero (Fig. 3a-d). As Figure 3 illustrates, most of the distributions were centered near zero, except for a slight but significant positive bias during delay 2 (p<0.005; sign test; Fig. 3d), indicating higher functional connectivity during the identity rule. In our second approach, we tested whether each functionally connected pair of units displayed a significant difference in r_sc_ between the two rules. The occurrence of significant differences (false discovery rate < 0.05, shuffled surrogates) were 0.9%, 0.8%, 1.0% and 1.1% for the pre-sample, sample, delay 1 and delay 2 epochs, respectively.

**Figure 3:**
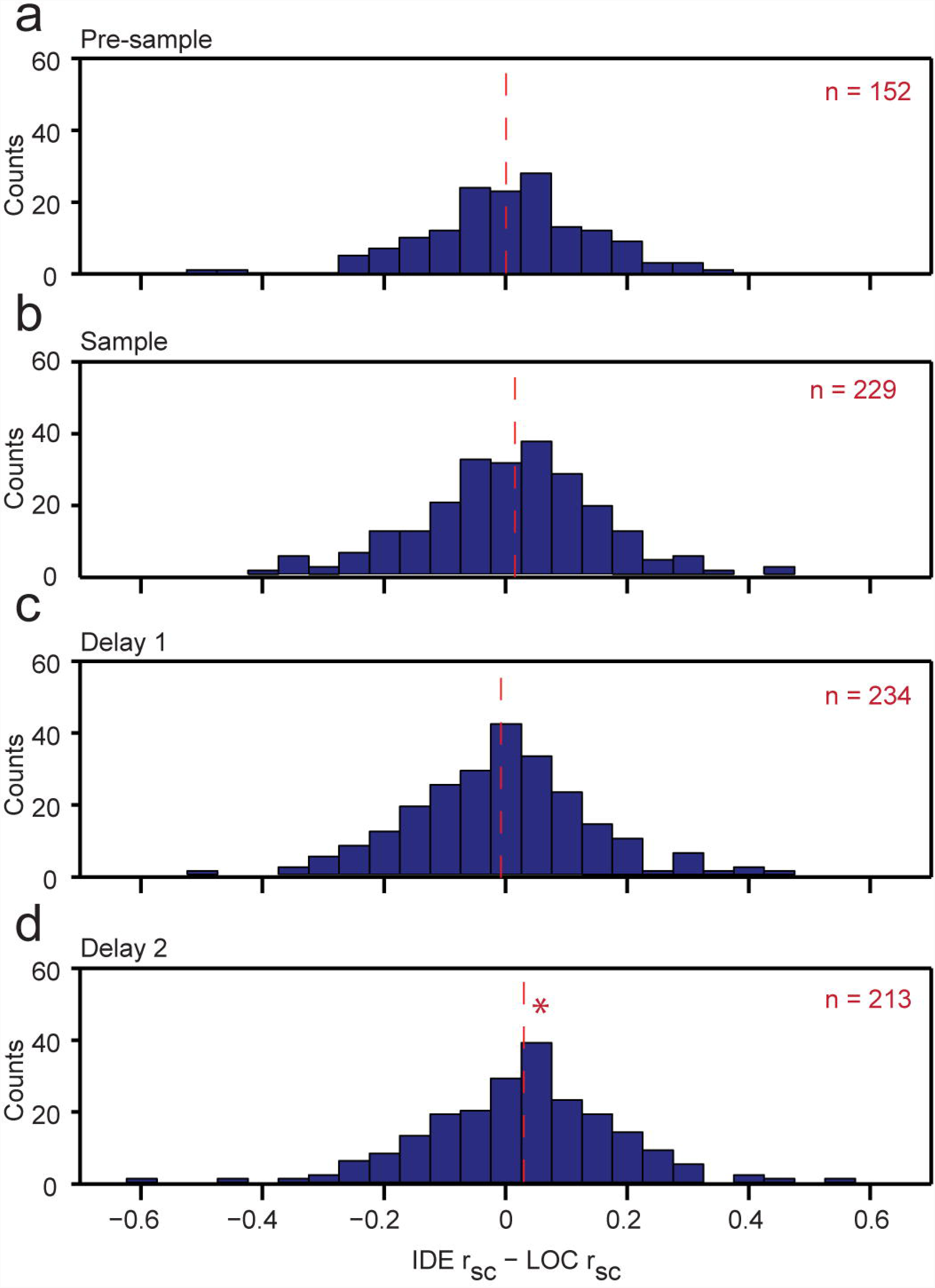
Rule preference of spike-count correlations. (a-d) Histograms of the differences in r_sc_ between the two rules for the pre-sample (a), sample (b), delay 1(c) and delay 2 (d) task periods. The median of each distribution is indicated by a dashed vertical line and is significantly different from zero only during delay 2 (p<0.005; Sign test).

Because r_sc_ is known to be sensitive to variations in firing rate (for a review see Cohen & Kohn, 2011), we compared the differences in r_sc_ to the differences in the geometric means of the firing rates between the two rules. This analysis revealed no relationship in any of the four epochs (Fig. S3), indicating that the result in Figure 3d is not readily accounted for by changes in firing rates.

To determine the anatomical distribution of functional connectivity during the task, we calculated the incidence of functionally connected units for each pair of cortical areas (Fig. 4). This revealed a widespread, but non-uniform, distribution of functional interactions among the areas sampled. The highest incidence occurred between units in the same cortical area (PE-PE, dPFC-dPFC, PEC-PEC). While the highest incidence of long-range prefrontal to posterior parietal functionally connected units occurred in PG-dPFC and LIP-dPFC. In general, we find a high incidence of dPFC-PG-LIP functional interactions (Fig. 4, black bars), similar to the results from the firing rate analysis.

**Figure 4:**
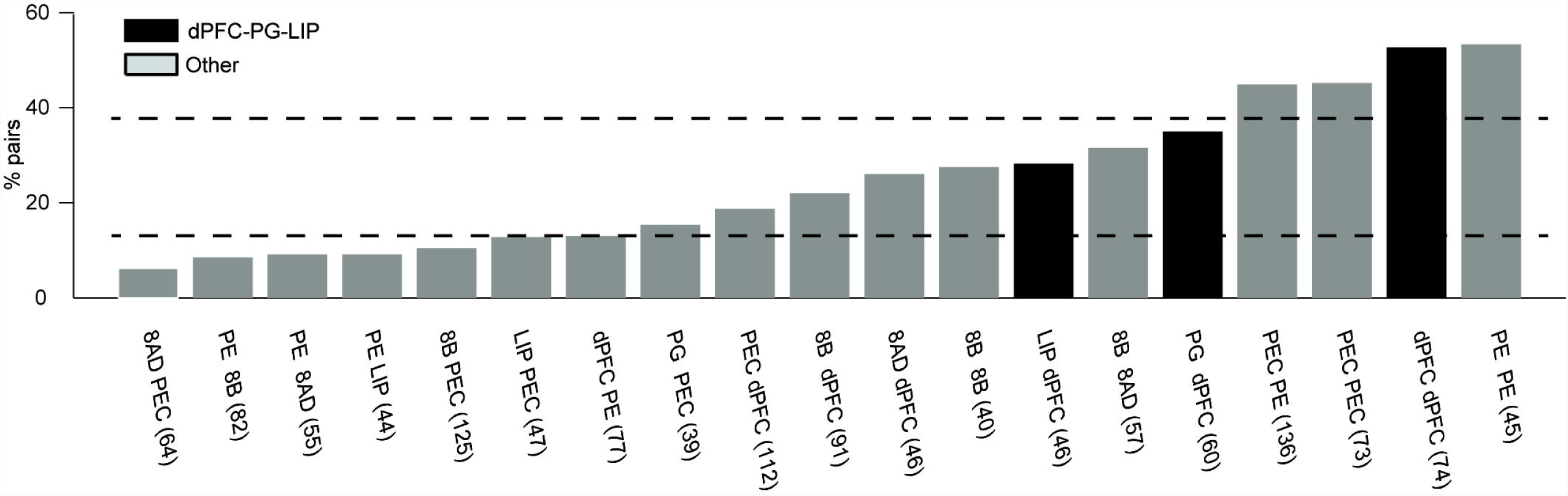
Incidence of significant spike-count correlations. A pair of units was considered significant if the spike-count correlation was significant during at least one period of the task for at least one of the rules. The numbers in parentheses on the x-axis indicate the sample size. Data are shown for pairs of cortical areas with a sample size of at least 30 pairs. The horizontal dashed red lines show the confidence intervals, 2.5% and 97.5%, for a random and uniform distribution. Black bars indicate the areal combinations composed of areas dPFC, PG, and LIP. All other areal combinations are in gray.

### Spike-Spike Coherence

As spike-count correlations may be driven by fluctuations operating at different frequencies, we further characterized the occurrence of trial-to-trial correlations in firing rates using multi-taper spectral coherence on the functionally connected units (Mitra and Pesaran, 1999; Bokil et al., 2007). This quantity estimates the frequency range of correlated fluctuations in activity between two units but does not necessarily imply the presence of oscillations. Consistent with a previous study (Mitchell et al., 2009), we found that correlated activity largely occurred at frequencies below 7 Hz. The highest incidence of significant coherence (p<0.01) occurred during delay 2 (Fig. 5). This result was still present even if the pairs of neurons were stratified with respect to their firing rates (Fig. S4). Altogether, this indicates that coherent spiking activity was enhanced in functionally connected units during the late phase of the memory delay.

**Figure 5:**
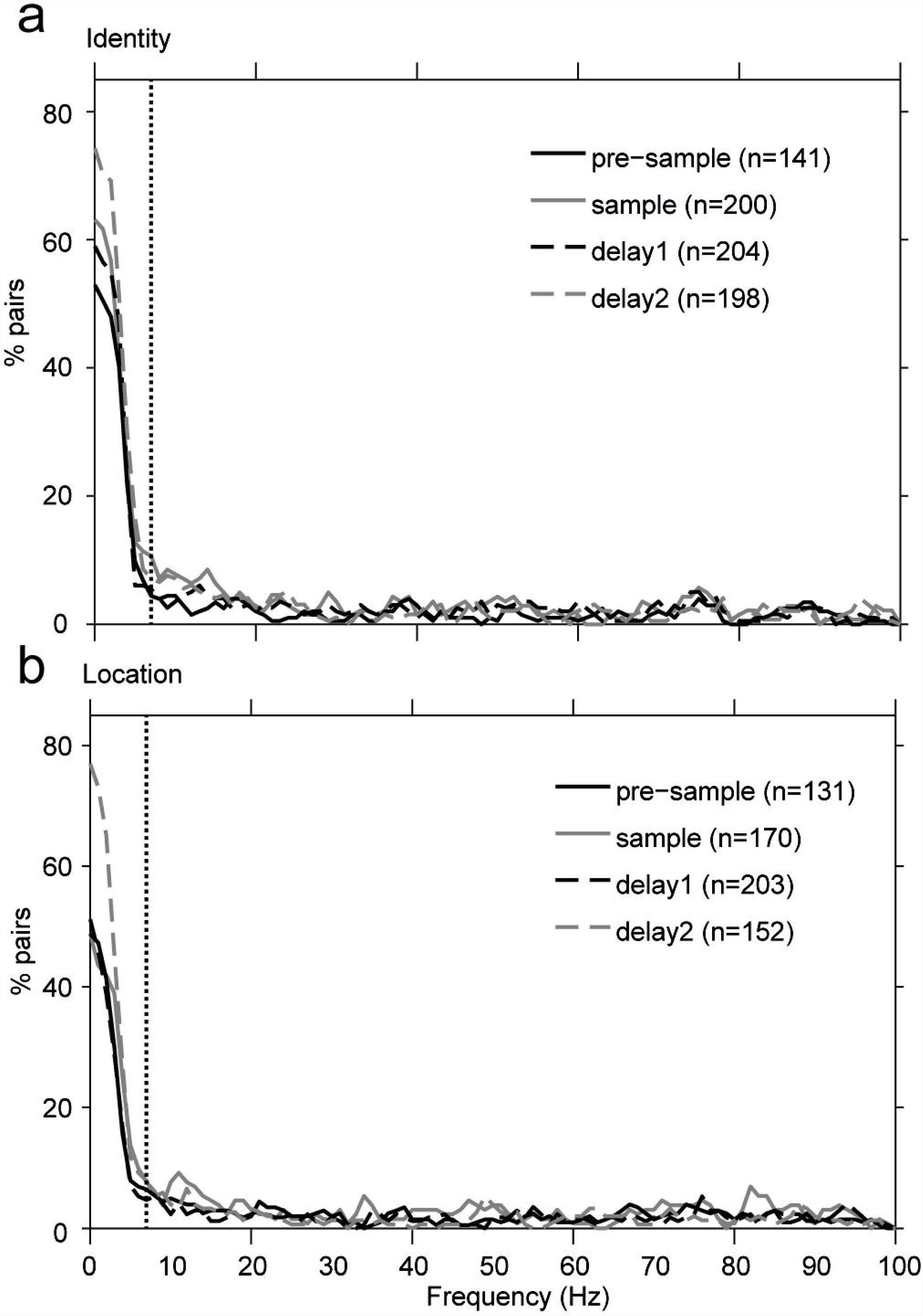
Analysis of the functionally connected units in the frequency domain. Coherence values were calculated for functionally connected units with significant (p<0.01; randomized surrogates) and positive r_sc_ during the identity (a) or the location rule (b). The dashed vertical lines are set at 7 Hz. The number of functionally connected units is displayed as a function of rule and epoch (n=sample size).

## DISCUSSION

During a delayed match-to-sample task involving a non-instructed switch between a location- and an identity-matching rule, many prefrontal and parietal units displayed rule selective activity throughout the task, even before the presentation of the sample stimulus. These rule-selective responses were most prevalent in the dorsal prefrontal cortex (dPFC) and in areas lateral to and within the lateral bank of the intra-parietal sulcus (PG and LIP). Positive spike count correlations between units were common, elevated with respect to the pre-sample period, and occurred primarily at low frequencies (< 7 Hz). Furthermore, these correlations were very prominent within dPFC and between dPFC and areas PG and LIP. Rule-selective functional connectivity was observed at the population level during the second half of the working memory delay (Fig. 3), but the incidence of individual pairs with significant rule-selectivity was low (< 5%) during all periods of the task.

Consistent with the role of prefrontal and posterior parietal areas in the active representation of behavioral rules (Asaad et al., 2000; Wallis et al., 2001; Wallis and Miller, 2003; Stoet and Snyder, 2004; Buckley et al., 2009; Gail et al., 2009; Goodwin et al., 2012), we observed rule selective activity during the sample and delay periods of the task. Interestingly, we also found that area PEC, which displays visual, somatosensory and motor responses (Batista et al., 1999; Battaglia-Mayer et al., 2001; Ferraina et al., 2001; Squatrito et al., 2001; Breveglieri et al., 2008), also demonstrated task dependent changes in rule selectivity. These results support the notion that rule processing involves a widespread network of cortical areas.

Our results also reveal the presence of rule information prior to the presentation of the sample stimulus. This result may initially seem odd. However, the matching rules were presented in blocks and once the behavioral performance reached 75%, the animal was probably aware of which rule was currently being enforced. A similar finding has been reported previously in prefrontal cortex (Asaad et al., 2000; Mansouri et al., 2006). Here, we extend this finding by showing that parietal areas also display rule specific activity prior to the sample stimulus. In fact, area PG showed a higher incidence of rule selectivity than dPFC (Fig. 1f).

It is well known that the responses of units to the repeated presentation of the same stimulus are highly variable. Population coding models consider this variability to be due to internal noise (Pouget et al., 2000). Across units, co-fluctuations in firing rates from trial-to-trial are thought to originate from common input and are similarly termed noise correlations (referred to as spike-count correlations in the current study). Under the population-coding model, changes in correlation affect the amount of information conveyed by the population (Averbeck et al., 2006). While the precise effects of increases, decreases and the sign of correlations on neuronal processing are not well understood (Averbeck et al., 2006), recent studies support the population-coding model by demonstrating that a decrease in correlation between nearby neurons is associated with selective attention (Cohen and Maunsell, 2009; Mitchell et al., 2009). In contrast, we find a marked increase in the incidence of significant positive correlations among units located in widely distributed cortical areas. This suggests that our results need to be interpreted differently. If we consider prefrontal and posterior parietal cortices as part of a widely interconnected network (Petrides and Pandya, 1984; Cavada and Goldman-Rakic, 1989a; Cavada and Goldman-Rakic, 1989b; Markov et al., 2014), and often co-activated during attention-demanding tasks (Goldman-Rakic, 1988; Kastner and Ungerleider, 2000; Salazar et al., 2012; Crowe et al., 2013; Dotson et al., 2014), then the resulting recurrent interactions would be expected to lead to an increase in functional connectivity (Heinzle et al., 2007) revealed by increases in spike count correlations (Pesaran et al., 2008; Fuster and Bressler, 2012).

Finally, we tested if rule guided working memory processes are correlated with changes in the functional connectivity between units. Based on the results from the firing rate analysis, we expected to find rule-dependent differences in r_sc_ during all periods of the task. However, we found only weak evidence for rule-dependent differences in the population and pair-wise analyses. At the population level, we found a slight but significant shift in magnitude toward the identity rule during the late delay period (Fig. 3). At the pairwise level, only a small percentage of unit pairs (< 5%) displayed changes in correlation magnitude with respect to the two rules. Altogether, these results suggest that changes in functional connectivity, as measured by spike count correlations, contribute only weakly to rule guided behavior.

There are several methodological considerations to take into account before interpreting spike-count correlations. First, spike-count correlations are sensitive to changes in firing rates. We show in Figure S3 that the changes, if any, in r_sc_ between rules are not readily explained by changes in the geometric mean firing rate. Also, the reported values of r_sc_ may appear to be high compared to values from previous studies. This may be due to the fact that most of the correlation estimates originated from multi-unit instead of single unit signals (Cohen and Kohn, 2011). Another methodological consideration is task design. Both the identity and location matching rules were performed during each recording session and the first matching rule was randomly chosen between the two rules. Unfortunately, it was rare for the animals to perform more than one rule switch, thereby limiting the number of replicated measures for each rule. As a consequence, we were unable to consistently control for the influence of other factors, such as fatigue or satiation that may change over the course of a session. This may have led us to underestimate the incidence of rule-dependent changes in r_sc_.

In conclusion, a convergence occurred in our data with respect to the areas showing the highest percentages of task and rule specificity (dPFC, PG, and LIP), supporting the previously mentioned evidence that dPFC and areas lateral to and within the lateral bank of the intraparietal sulcus play a major role in both working memory and rule-guided behavior. However, we also demonstrate that several areas, not previously considered under these conditions, also exhibit task and rule related changes in activity. These findings suggest that rule guided visual working memory is supported by several functionally connected areas that are widely distributed across the brain.

## Supporting information

## ACKNOLWDEGEMENTS

This work was supported by grants from the National Institute of Mental Health (MH069374, MH081162), the National Institute of Neurological Disorders and Stroke (NS059312), a Swiss National Science Foundation award and an award from the Kopriva Foundation. We thank Behrad Noudoost, Kelsey Clark, Steven Hoffman, Tiphani Lynn and Rebecca Brooker for comments on previous versions of the manuscript. The author declares no competing financial interests.

