## Supplementary material for "Widespread changes in firing rate and functional connectivity across the fronto-parietal network during rule guided working memory"

### Figure Legends

Figure S1. Representative example of the shuffle-fitting procedure for p-values estimates of cross-correlations. A histogram (100 equivalent bins, blue bars) of the surrogate r_sc_ population (500 iterations) was generated and a Generalized Extreme Value distribution was fit to this population (violet line). This fitted function was then used to estimate the p-value of the corresponding r_sc_ (red star).

Figure S2. Distribution of firing rates as a function of cortical area. The firing rates were calculated separately for the identity (a) and location (b) rules during the memory delay.

Figure S3. Rule differences in firing rates and in spike-count correlations. The panels show scatter plots of the rule difference (IDENTITY minus LOCATION) in r_sc_ as a function of the rule difference (IDENTITY minus LOCATION) in the averaged firing rate (geometric mean) between the participating neurons. Each panel corresponds to a different epoch of the task (from left to right: presample, sample, delay 1 and delay 2).

Figure S4. Relationship between firing rate and coherent spiking. Coherence values were calculated for functionally connected units with significant (p<0.01; randomized surrogates) and positive r_sc_ during the identity (a) or the location rule (b). The data were split according to the median firing rate of the two participating neurons during the delay period (800 msec). The pairs with low (0 to 33.3^th^ percentiles), medium (33.3^th^ to 66.6^th^ percentiles) and high firing rates (66.6^th^ to 100^th^ percentiles) are displayed respectively from left to right and for each epoch separately (colored lines). In all cases, the highest incidence of significant coherence occurred during delay 2 in the low frequencies. The blue lines are set at 7 Hz.
