## Supplementary figures and images for "Widespread changes in firing rate and functional connectivity across the fronto-parietal network during rule guided working memory"

### Supplementary file 2

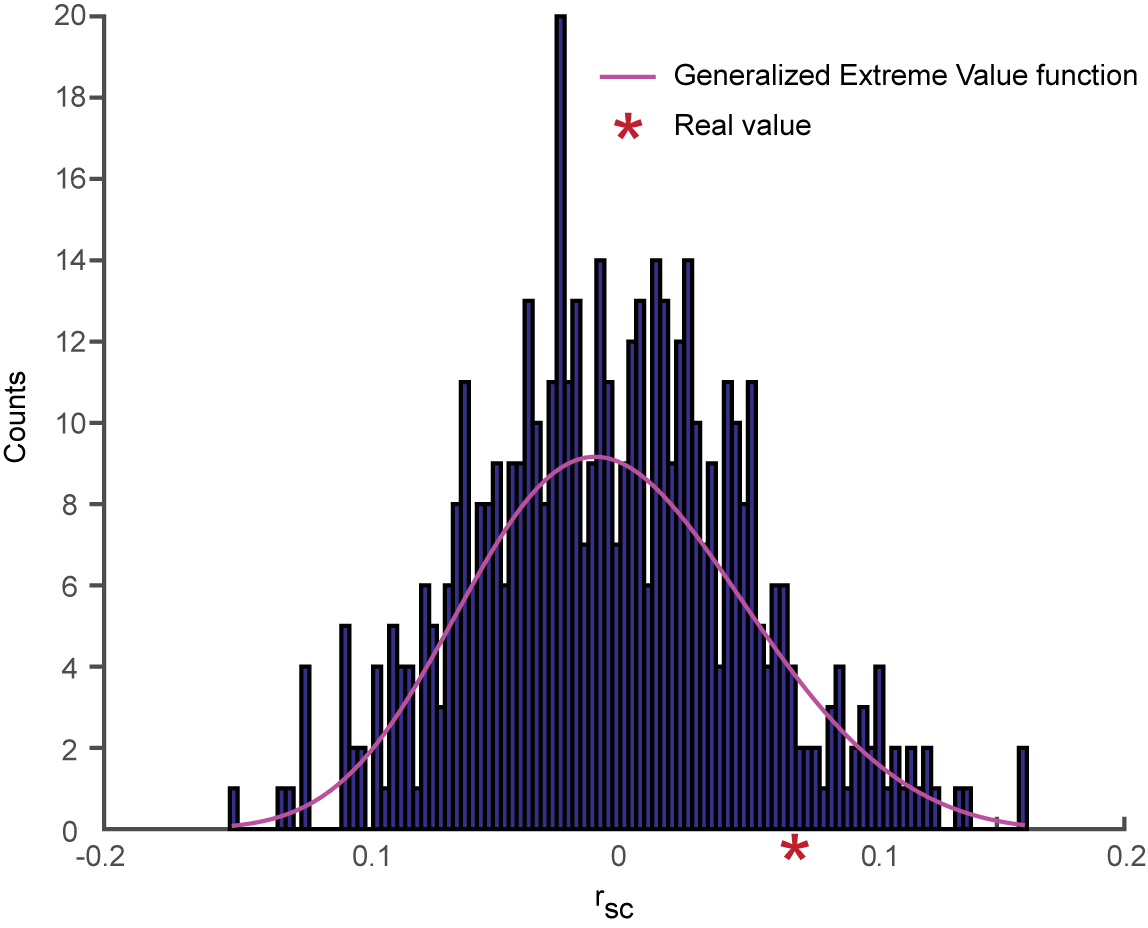

### Supplementary file 3

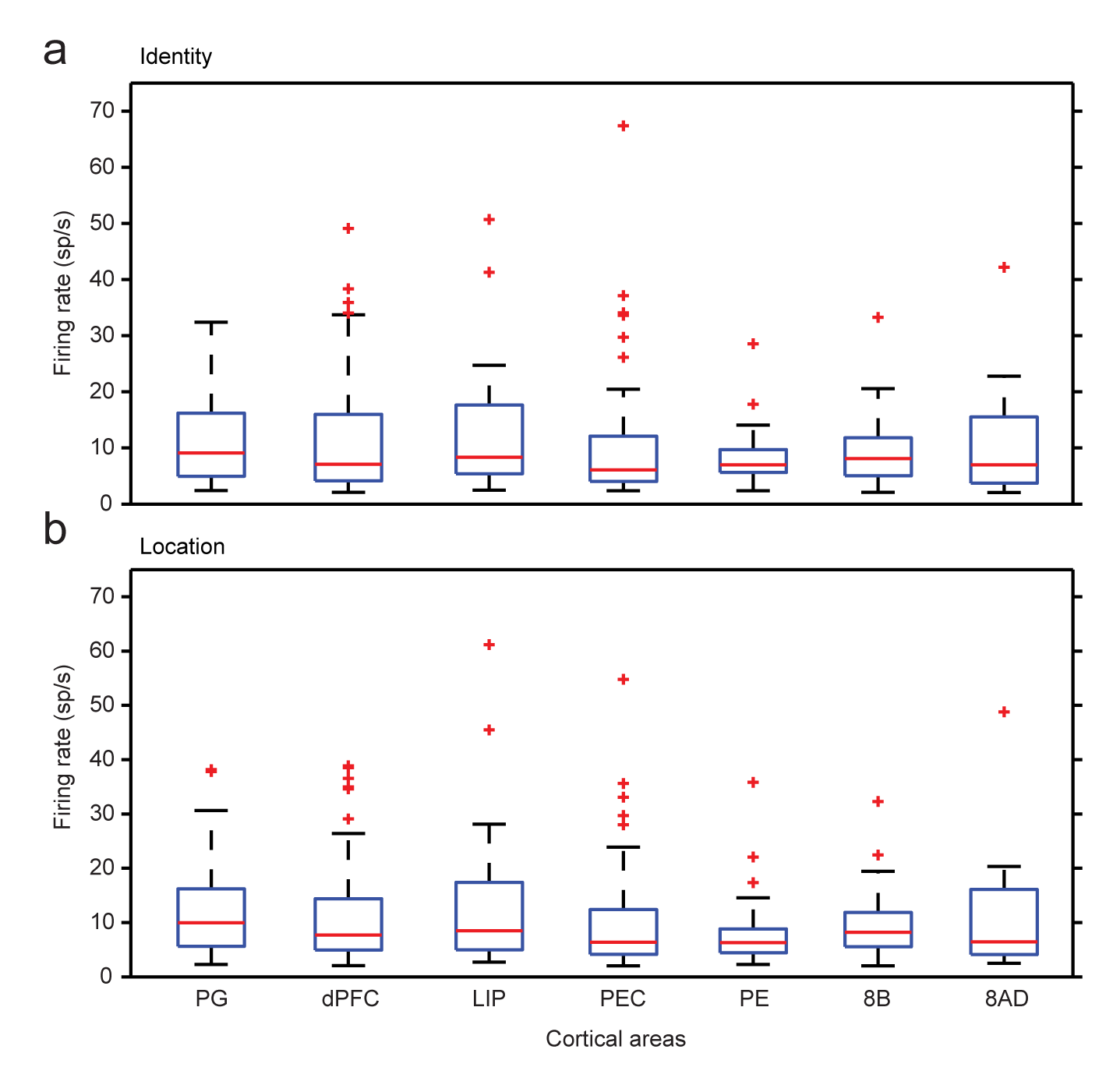

### Supplementary file 4

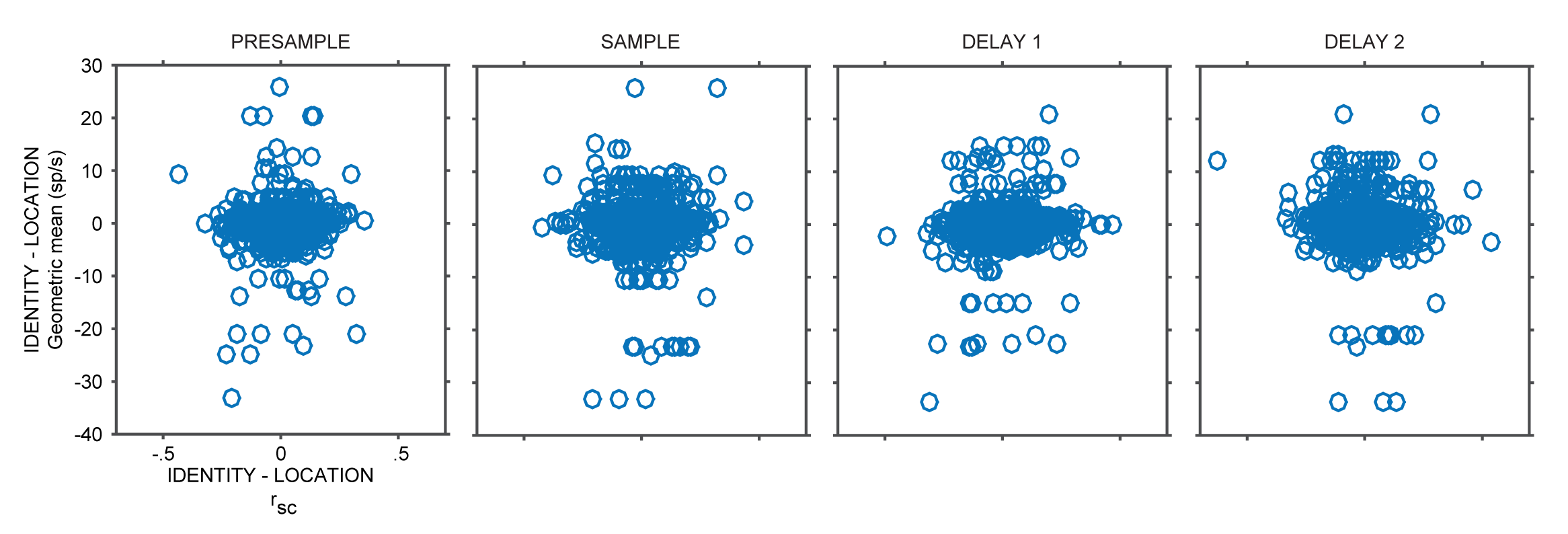

### Supplementary file 5

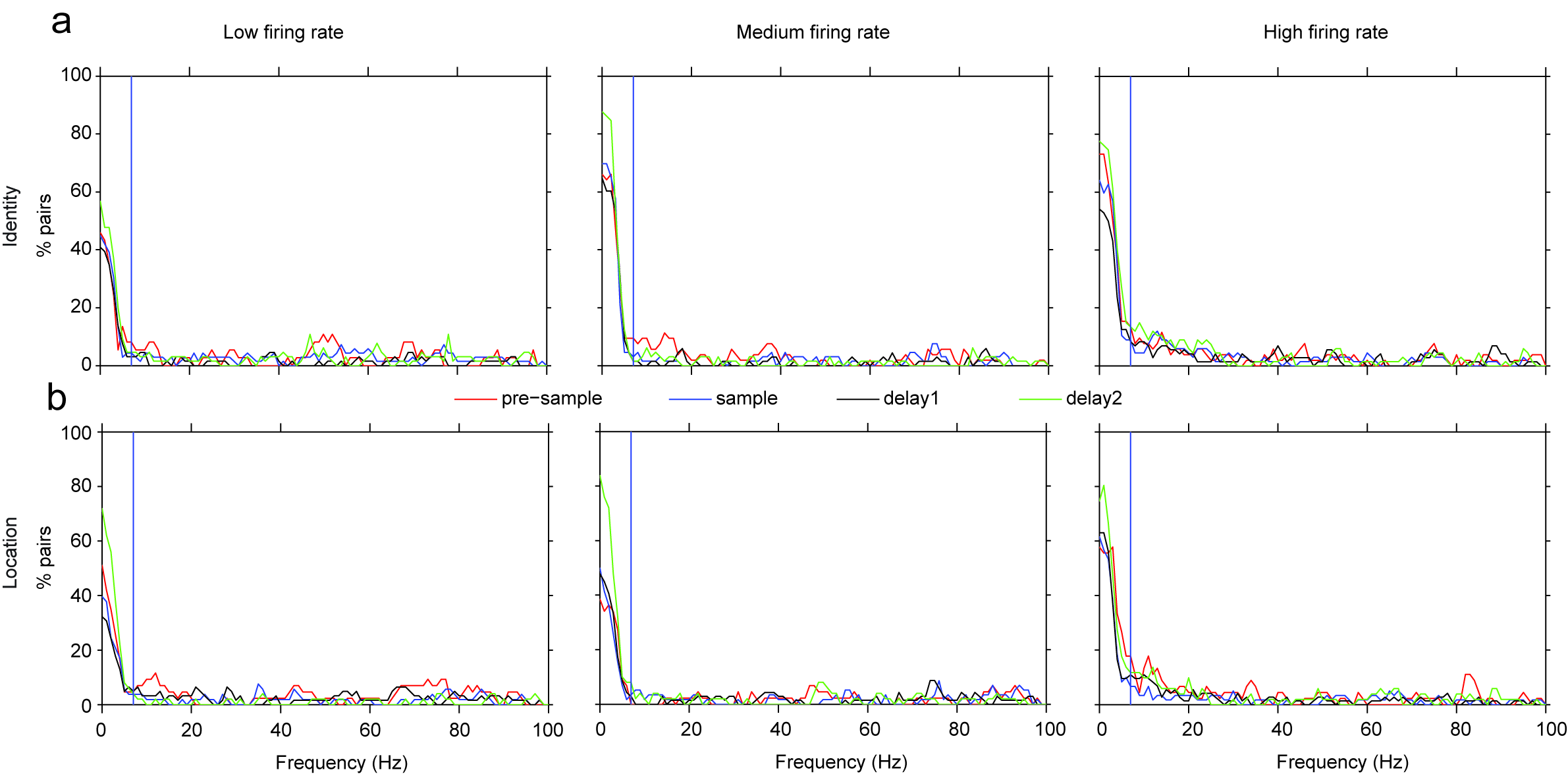
